# DNA profiling reveals *Neobenedenia girellae* as the primary culprit in global fisheries and aquaculture

**DOI:** 10.1101/304261

**Authors:** Alexander K. Brazenor, Terry Bertozzi, Terrence L. Miller, Ian D. Whittington, Kate S. Hutson

**Affiliations:** Marine Parasitology Laboratory, Centre for Sustainable Tropical Fisheries and Aquaculture and the College of Marine and Environmental Sciences, James Cook University, Townsville, Queensland 4811, Australia.; The South Australian Museum, North Terrace, Adelaide, SA 5000, Australia.; School of Biological Sciences, University of Adelaide, Adelaide, SA, 5005, Australia.; Centre for Sustainable Tropical Fisheries and Aquaculture and the College of Marine and Environmental Sciences, James Cook University, Cairns, Queensland 4870, Australia.; Fish Health Laboratory, Department of Fisheries Western Australia, Perth, Western Australia 6151, Australia.

**Keywords:** phylogeny, host specificity, Monogenea, Capsalidae, skin fluke, *Neobenedenia melleni*, cryptic

## Abstract

Accurate identification of parasite species and strains is crucial to mitigate the risk of epidemics and emerging disease. Species of *Neobenedenia* are harmful monogenean ectoparasites that infect economically important bony fishes in aquaculture worldwide, however, the species boundaries between two of the most notorious taxa, *N. melleni* and *N. girellae*, has been a topic of contention for decades. Historically, identifications of *Neobenedenia* isolates have overwhelmingly been attributed to *N. melleni*, and it has been proposed that *N. girellae* is synonymous with *N. melleni*. We collected 33 *Neobenedenia* isolates from 22 host species spanning nine countries and amplified three genes including two nuclear (*Histone 3* and *28S rDNA*) and one mitochondrial (*cytochrome b*). Four major clades were identified using Maximum Likelihood and Bayesian inference analyses; clades A-D corresponding to *N. girellae, N. melleni, N. longiprostata* and *N. pacifica* respectively. All unidentified isolates and the majority of *Neobenedenia* sequences from GenBank fell into clade A. The results of this study indicate that *N. girellae* is a separate species to *N. melleni*, and that a large proportion of previous samples identified as *N. melleni* may be erroneous and a revision of identifications is needed.

The large diversity of host species that *N. girellae* is able to infect as determined in this study and the geographic range in which it is present (23.8426° S and 24.1426° N) makes it a globally cosmopolitan species and a threat to aquaculture industries around the world.

## 1. Introduction

The Monogenea, one of three predominantly parasitic classes in the phylum Platyhelminthes, predominantly comprises ectoparasitic flukes that infect the gills and skin surfaces of marine and freshwater fishes (Whittington, 2004). Monogeneans in the family Capsalidae (Monopisthocotylea) are recognised as virulent pathogens of finfish in sea-cage and semi-open pond aquaculture and are able to multiply rapidly in high-density aquaculture environments as they have a direct, single host life cycle (Jahn and Kuhn, 1932; Ogawa, 1996). Some species are responsible for considerable epidemics (Bauer and Hoffman, 1976; Deveney et al., 2001; Paperna and Overstreet, 1981; Whittington et al., 2004). According to current taxonomic classification, the Capsalidae comprises nine subfamilies, 57 genera, and over 300 species, most of which are ectoparasitic on marine fishes (Gibson et al., 2010). One of the most notorious of capsalid genera is *Neobenedenia*, Yamaguti 1963, which have a large host range and have been implicated in causing severe pathology and devastating economic losses in aquaculture worldwide.

*Neobenedenia* spp. harm fish by mechanical attachment and subsequent grazing on the epithelial surface of their host, which can cause epidermal degradation, dermal erosion, inflammation and allow the ingress of secondary pathogens (Kaneko et al., 1988; Trujillo-González et al., 2015). *Neobenedenia* currently comprises six species and has gained global attention primarily due to the notoriety of the type species, *Neobenedenia melleni*, MacCallum 1927, which was originally described from tropical fishes held in the New York Aquarium (MacCallum, 1927; Whittington and Horton, 1996). While the origin of the infection is not clear, it is thought that the parasite may have been introduced from the Caribbean through the ornamental fish trade (Dyer et al., 1992; Whittington and Horton, 1996). *Neobenedenia melleni* is infamous as a widespread pathogen in aquaria and aquaculture and the list of recorded host taxa spans over 100 species from 30 families and five orders of teleosts (Bullard et al., 2000; Harris et al., 2004; Whittington and Horton, 1996; Whittington, 2004; Whittington et al., 2004). The broad host specificity reported for *N. melleni* is atypical for monogeneans, where it is considered that approximately 80% of all monogenean species only infect one host species and usually only in a single ocean (Byrnes and Rohde, 1992; Rohde, 1979; Whittington, 1998; Whittington et al., 2000).

Given their specialised lifestyle, it is not surprising that the morphology of species of *Neobenedenia* is highly conserved and accurate identification has proven challenging (Ogawa et al., 1995; Whittington, 2004; Whittington et al., 2004). Of particular controversy and confusion has been the delineation of *N. melleni* and *Neobenedenia girellae*, Hargis, 1955. *Neobenedenia girellae* was originally described by Hargis (1955) infecting opaleye, *Girella nigricans*, Ayres 1860, from California. Twenty-six additional host species have been subsequently reported from a range of localities (Nigrelli, 1947; Bravo-Hollis, 1958; Bravo-Hollis and Deloya, 1973; Gaida and Frost, 1991; Love et al., 1984; Moser and Haldorson, 1982; Ogawa et al., 1995). Twenty years ago, Whittington and Horton (1996) synonymised *N. girellae* with *N. melleni* based on morphology and host specificity. This decision was not unanimously accepted and many authors continue to use ‘*N. girellae*’ in the scientific literature (e.g. Koesharyani et al., 1999; Ogawa et al., 2006; Ogawa and Yokoyama, 1998; Wang et al., 2004; Yoshinaga et al., 2000; Zhang et al., 2014).

Studies employing molecular techniques in systematics have revealed a large number of morphologically similar parasite species that were previously recognized as a single taxon but are actually genetically distinct (Blasco-Costa et al., 2010; Donald et al., 2004; Freeman & Ogawa, 2010; Leung et al., 2009; Miura et al. 2005; Saijuntha et al., 2007; Sepúlveda and González, 2014; Wu et al., 2005; Xiao et al. 2005). For example, Sepúlveda and González (2014) found that a related capsalid species, *Benedenia seriolae*, Yamaguti 1934, is in fact a complex of cryptic species and not a single taxon. The first molecular-based evidence suggesting that a species complex may be present among *Neobenedenia* spp. was presented by Whittington et al. (2004) where *28S rDNA* sequences were compared between two *Neobenedenia* isolates identified as *N. melleni*. This study indicated that distinct taxa may be present within a complex of morphologically similar individuals. Since this research, very few molecular studies have addressed *Neobenedenia* taxa in order to resolve the confusion. A brief study comparing *N. melleni* and *N. girellae* by Wang et al. (2004) used short sequences of *28S rDNA* and indicated that the synonymy of these species proposed by Whittington and Horton (1996) was supported. However, a more comprehensive study by Perkins et al. (2009), which addressed the phylogeny of many capsalid species using multiple genes (*28S rDNA, Histone 3*, and *Elongation Factor* 1α), seems to support the opposite view; that *N. melleni* and *N. girellae* are two separate species.

*Neobenedenia* was first documented in Australian waters in 2000. An outbreak of *N. melleni* was reported in farmed barramundi, *Lates calcarifer*, Bloch 1790, and resulted in the death of over 200,000 fish (Deveney et al., 2001). However, it was only in 2011 that research on *Neobenedenia* in Australia began to develop following the collection of *Neobenedenia* sp. individuals from a fish farm in north Queensland and the subsequent establishment and maintenance of a continuous *in vivo* culture in the Marine Parasitology Laboratory at James Cook University, Australia (Hutson et al., 2012). Morphological identification of the species of *Neobenedenia* in culture has been problematic and all previous research from this laboratory has referred to the parasite as *Neobenedenia* sp. (e.g. Dinh Hoai and Hutson, 2014; Hutson et al., 2012; Trujillo-González et al., 2015a,b). There is a need to accurately identify the species of *Neobenedenia* currently being used as a parasite model in this laboratory so that research can be ascribed to the correct taxon allowing for more meaningful application.

The aim of this study was to use molecular characterisation methods to generate a robust diagnostic and phylogenetic framework for the identification of *Neobenedenia* spp. to underpin our understanding of the ecology of the taxa implicated in causing damage to wild and aquaculture fish stocks worldwide.

## 2. Materials and Methods

### 2.1. Sample collection

*Neobenedenia* specimens were collected between 2000 and 2015 from wild and captive fish in nine countries. The majority of Australian isolates were collected by the authors while many of the international samples were donated by research colleagues (Table 1). Samples were collected by removing live parasites from the skin of their hosts using a scalpel blade or after bathing the host in freshwater, which kills the parasites. Parasites were fixed and stored in 70% analytical grade ethanol.

**Table 1:**
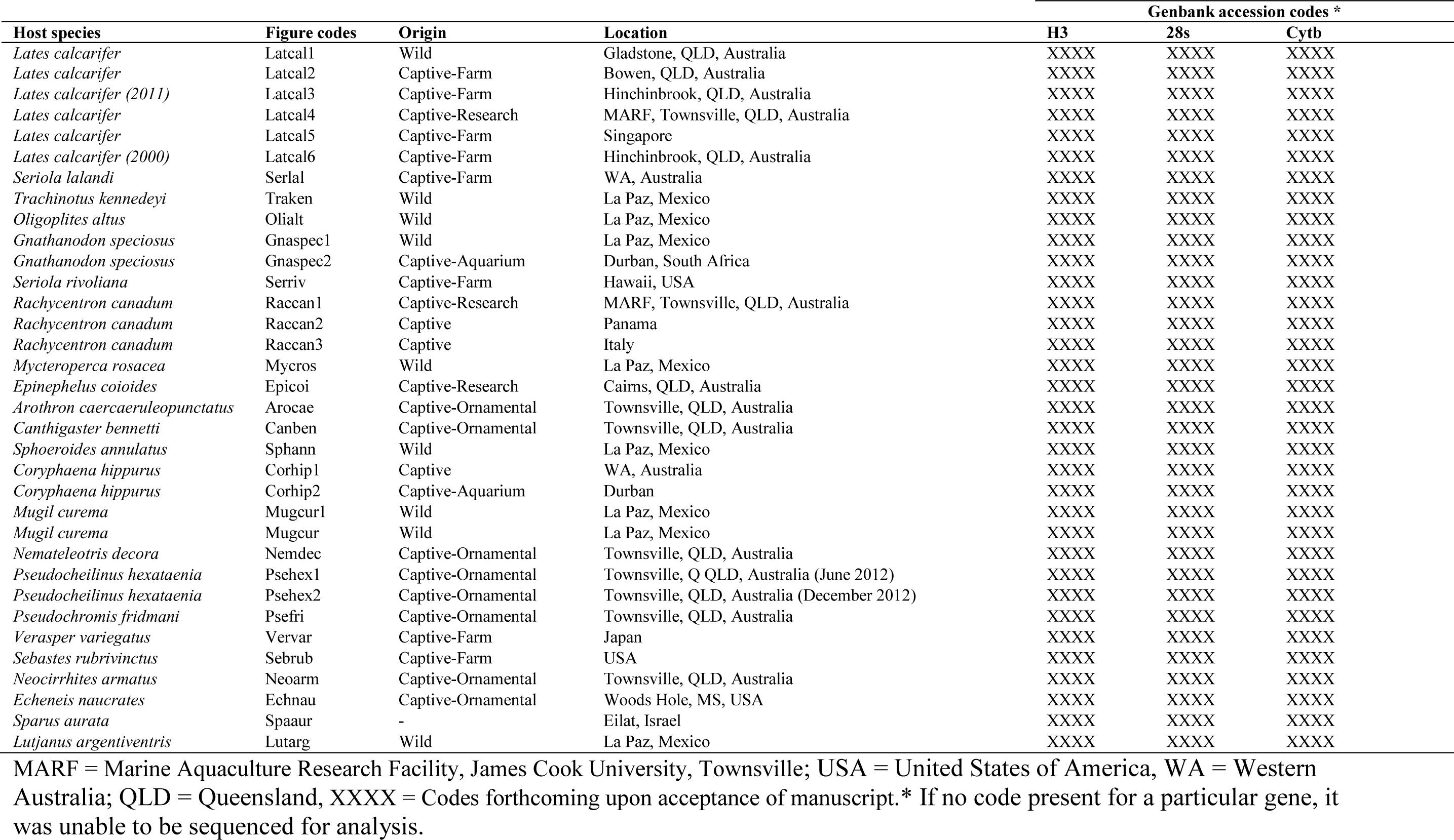
Host fish species from which *Neobenedenia* were sampled and used for sequencing from Australia and overseas.

| Host species | Figure codes | Origin | Location | Genbank accession codes * |  |  |
| --- | --- | --- | --- | --- | --- | --- |
|  |  |  |  | H3 | 28s | Cytb |
| *Lates calcarifer* | Latcal1 | Wild | Gladstone, QLD, Australia | XXXX | XXXX | XXXX |
| *Lates calcarifer* | Latcal2 | Captive-Farm | Bowen, QLD, Australia | XXXX | XXXX | XXXX |
| *Lates calcarifer* (2011) | Latcal3 | Captive-Farm | Hinchinbrook, QLD, Australia | XXXX | XXXX | XXXX |
| *Lates calcarifer* | Latcal4 | Captive-Research | MARF, Townsville, QLD, Australia | XXXX | XXXX | XXXX |
| *Lates calcarifer* | Latcal5 | Captive-Farm | Singapore | XXXX | XXXX | XXXX |
| *Lates calcarifer* (2000) | Latcal6 | Captive-Farm | Hinchinbrook, QLD, Australia | XXXX | XXXX | XXXX |
| *Seriola lalandi* | Serlal | Captive-Farm | WA, Australia | XXXX | XXXX | XXXX |
| *Trachinotus kennedeyi* | Traken | Wild | La Paz, Mexico | XXXX | XXXX | XXXX |
| *Oligoplites altus* | Olialt | Wild | La Paz, Mexico | XXXX | XXXX | XXXX |
| *Gnathanodon speciosus* | Gnaspec1 | Wild | La Paz, Mexico | XXXX | XXXX | XXXX |
| *Gnathanodon speciosus* | Gnaspec2 | Captive-Aquarium | Durban, South Africa | XXXX | XXXX | XXXX |
| *Seriola rivoliana* | Serriv | Captive-Farm | Hawaii, USA | XXXX | XXXX | XXXX |
| *Rachycentron canadum* | Raccan1 | Captive-Research | MARF, Townsville, QLD, Australia | XXXX | XXXX | XXXX |
| *Rachycentron canadum* | Raccan2 | Captive | Panama | XXXX | XXXX | XXXX |
| *Rachycentron canadum* | Raccan3 | Captive | Italy | XXXX | XXXX | XXXX |
| *Mycteroperca rosacea* | Mycros | Wild | La Paz, Mexico | XXXX | XXXX | XXXX |
| *Epinephelus coioides* | Epicoi | Captive-Research | Cairns, QLD, Australia | XXXX | XXXX | XXXX |
| *Arothron caeruleopunctatus* | Arocae | Captive-Ornamental | Townsville, QLD, Australia | XXXX | XXXX | XXXX |
| *Canthigaster bennetti* | Canben | Captive-Ornamental | Townsville, QLD, Australia | XXXX | XXXX | XXXX |
| *Sphoeroides annulatus* | Sphann | Wild | La Paz, Mexico | XXXX | XXXX | XXXX |
| *Coryphaena hippurus* | Corhip1 | Captive | WA, Australia | XXXX | XXXX | XXXX |
| *Coryphaena hippurus* | Corhip2 | Captive-Aquarium | Durban | XXXX | XXXX | XXXX |
| *Mugil curema* | Mugcur1 | Wild | La Paz, Mexico | XXXX | XXXX | XXXX |
| *Mugil curema* | Mugcur | Wild | La Paz, Mexico | XXXX | XXXX | XXXX |
| *Nemateleotris decora* | Nemdec | Captive-Ornamental | Townsville, QLD, Australia | XXXX | XXXX | XXXX |
| *Pseudocheilinus hexataenia* | Psehex1 | Captive-Ornamental | Townsville, Q QLD, Australia (June 2012) | XXXX | XXXX | XXXX |
| *Pseudocheilinus hexataenia* | Psehex2 | Captive-Ornamental | Townsville, QLD, Australia (December 2012) | XXXX | XXXX | XXXX |
| *Pseudochromis fridmani* | Psefri | Captive-Ornamental | Townsville, QLD, Australia | XXXX | XXXX | XXXX |
| *Verasper variegatus* | Vervar | Captive-Farm | Japan | XXXX | XXXX | XXXX |
| *Sebastes rubrivinctus* | Sebrub | Captive-Farm | USA | XXXX | XXXX | XXXX |
| *Neocirrhites armatus* | Neoarm | Captive-Ornamental | Townsville, QLD, Australia | XXXX | XXXX | XXXX |
| *Echeneis naucrates* | Echnau | Captive-Ornamental | Woods Hole, MS, USA | XXXX | XXXX | XXXX |
| *Sparus aurata* | Spaaur | - | Eilat, Israel | XXXX | XXXX | XXXX |
| *Lutjanus argentiventris* | Lutarg | Wild | La Paz, Mexico | XXXX | XXXX | XXXX |

MARF = Marine Aquaculture Research Facility, James Cook University, Townsville; USA = United States of America, WA = Western Australia; QLD = Queensland, XXXX = Codes forthcoming upon acceptance of manuscript.\* If no code present for a particular gene, it was unable to be sequenced for analysis.

A slice of tissue (sliver) was removed from the right-hand side margin of individual parasites, opposite the reproductive organs, taking care to avoid the gut. The remainder of the specimen was washed three times using distilled H_2_O and stained with Haematoxylin. Parasites were then dehydrated through an alcohol series, cleared in methyl salicylate or cedarwood oil, and mounted on microscope slides in Canada balsam for further study (Hutson et al., 2012). At least one mounted specimen collected per host by A.K. Brazenor and K.S. Hutson was accessioned into the Queensland Museum helminthology collection. Specimens from the late I.D. Whittington’s collection were accessioned to the Australian Helminthological Collection at the South Australian Museum (Table 1).

### 2.2. DNA preparation, PCR amplification, and amplicon sequencing

DNA was extracted from parasite slivers using either the PUREGENE DNA purification system (Gentra Systems) protocol for DNA purification from solid tissue or a QIAgen DNeasy Kit (QIAGEN Inc., Valencia, California, USA) according to the manufacturers’ protocols. PCR amplifications of partial *H3, 28S rDNA*, and *Cytb* sequence were carried out in 25μl reactions using the primers listed in Table 2 and either Amplitaq Gold DNA polymerase, or Phusion high-fidelity polymerase with the following reaction and cycling conditions: Amplitaq Gold – we followed Perkins et al. (2009) except that we standardised the annealing temperature at 55°C and used a maximum of 34 PCR cycles: Phusion - we used a final concentration of 5μL of 5xPhusion^®^ HF buffer, 0.5 μL of 10mM dNTPs, 1.25 μL of each primer (10mM), 0.25 μL of Phusion^®^ Tag DNA polymerase, and 4 μL of DNA template with an initial denaturation step of 98 °C for 30 s, followed by 35 cycles of PCR; denaturation at 98 °C for 10 s, annealing 53-62 °C for 20 s, extension at 72 °C for 30 s, with an additional final extension at 72 °C for 7.5 min. The double-stranded amplification products were visualised on 1.5% agarose gels and purified using a Multiscreen – PCR Plate (Millipore Corporation). Purified products were sent to the Australian Genome Research Facility for cycle-sequencing in both directions using the BigDye Terminator v3.1 cycle-sequencing kit (Applied Biosystems) on an AB3730xl capillary sequencer.

**Table 2:**
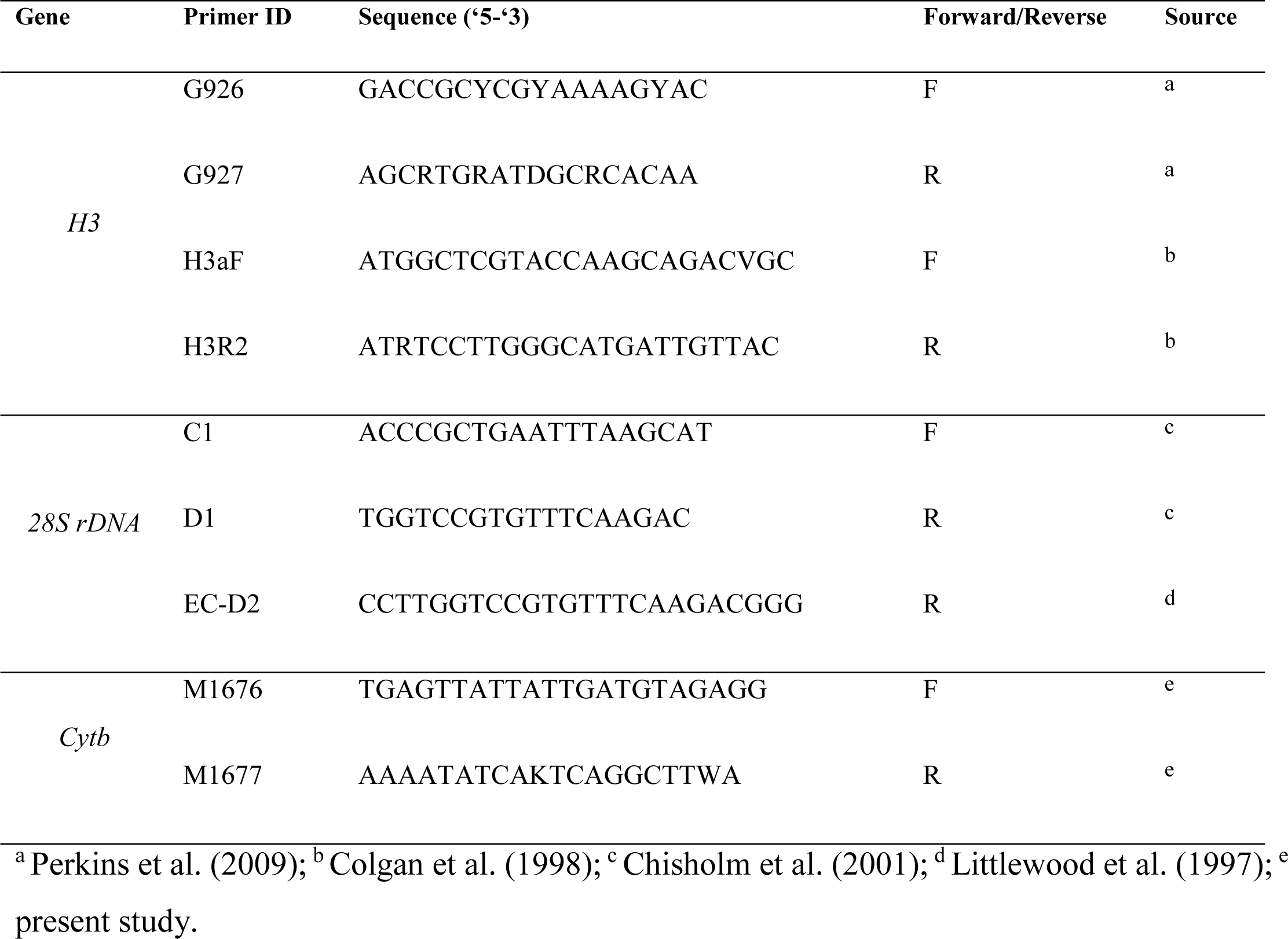
Primers used for PCR amplification of *Neobenedenia* spp. isolates.

### 2.3. Phylogenetic analyses

Sequence chromatograms were edited with SeqEd 1.0.3 (Applied Biosystems) and aligned using Se-Al2.0a11 (http://tree.bio.ed.ac.uk/software/seal/) using inferred amino acid translations (*H3* and *Cytb*) and the predicted *28S rDNA* secondary structure model for *Gyrodactylus salaris*, Malmberg 1957, (Matejusová and Cunningham, 2004). Subsequently, the *28S rDNA* alignment was trimmed to 411bp to remove sequences that could not be aligned unambiguously and sequences from all three genes concatenated to give a final alignment length of 1409bp. We were unable to amplify all target fragments for some samples and these are coded in the final alignment as “missing data”. All gene sequences have been deposited in GenBank (codes to be added following acceptance of manuscript). PartitionFinder v1.1.1 (Lanfear et al., 2012) was used to partition the data and select the most optimal model of nucleotide substitution for each partition based on the Akaike Information Criterion.

Bayesian phylogenetic analyses were run using the MPI version of MrBayes 3.2.6 (Ronquist et al., 2012) on a 12-core virtual machine on the NeCTAR research cloud under an Ubuntu 16.04 LTS image using Open MPI version 2.0.1 https://www.open-mpi.org/. Analyses employed two runs, each with four chains, with ten million steps, sampling every 1000 steps.

The first 20% of sampled topologies was discarded as burn-in based on stability of log likelihood values and that sampled topologies were essentially identical across runs, with standard deviation of split frequencies ~0.01 or less. Samples for numerical parameters were also essentially identical, with variance between versus within runs approaching unity (Ronquist et al., 2012).

The majority-rule consensus tree was constructed from the combined post-burn-in samples. Maximum likelihood (ML) analyses was conducted using the RAxML BlackBox server (http://embnet.vital-it.ch/raxml-bb/) implementing the methods of Stamatakis et al. (2008). Data were partitioned as recommended by PartitionFinder and run using the Gamma model of rate heterogeneity.

All *Neobenedenia Cytb* and *28S rRNA* sequences on GenBank were downloaded and aligned to our data. We also included the *Cytb* sequence of the only full mitochondrial genome of this genus (JQ038228). A neighbor-joining analysis of the aligned data for each locus was conducted in MEGA 6.06 (Tamura et al., 2013), using uncorrected *P* distance, in an attempt to reconcile the GenBank sequences with our phylogenetic framework.

## 3. Results

### 3.1. DNA sequence characteristics

Amplified *Cytb* and *H3* sequences did not contain any premature stop codons or frameshift mutations, contributing 704 and 292 characters respectively to the overall concatenated alignment. Even though we were able to amplify over 800bp of sequence for *28S rDNA*, the sequence spans a highly variable region which we were unable to unambiguously align, even when using the secondary structure of the *28S rDNA* sequence for *G. salaris* as a guide. Thus, we were forced to trim our *28S rDNA* sequences to 411 characters. Some *28S rDNA* and *H3* sequence chromatograms contained overlapping peaks indicative of heterozygous alleles and these sites were scored with IUPAC ambiguity codes for dimorphic sites. All individuals included in the analyses are represented by sequences of at least two of the genes (*Cytb* and *H3* or *28S rDNA*).

### 3.2. Phylogenetic analyses

Analysis of the concatenated dataset in PartitionFinder v1.1.1 suggested that three partitions (*Cytb* codon position 3; *Cytb* codon positions 1 and 2; *H3* codon positions 1, 2 and 3 and *28S rDNA*) were optimal for the data. PartitionFinder selected the General Time Reversible (GTR) model with a gamma distribution for rates across sites (GTR + G), GTR + G incorporating a proportion of invariable sites (GTR + I + G) and the Symmetrical model with a gamma distribution for rates across sites (SYM + G) for the partitions respectively.

Bayesian (Fig. 1) and Maximum Likelihood (ML) (Fig. 2) analyses yielded essentially identical topologies, with *Neobenedenia* samples forming a monophyletic clade to the exclusion of the outgroups and separated into four major clades (A-D). The *Neobenedenia* isolate recovered from *Lutjanus argentiventris*, Peters 1869, was the exception and was recovered as part of clade A in the Bayesian analysis (Fig. 1) but as a sister to clade A in the ML analysis (Fig. 2; labelled A* for clarity) albeit with low bootstrap support (BS 24%).

**Fig. 1.**
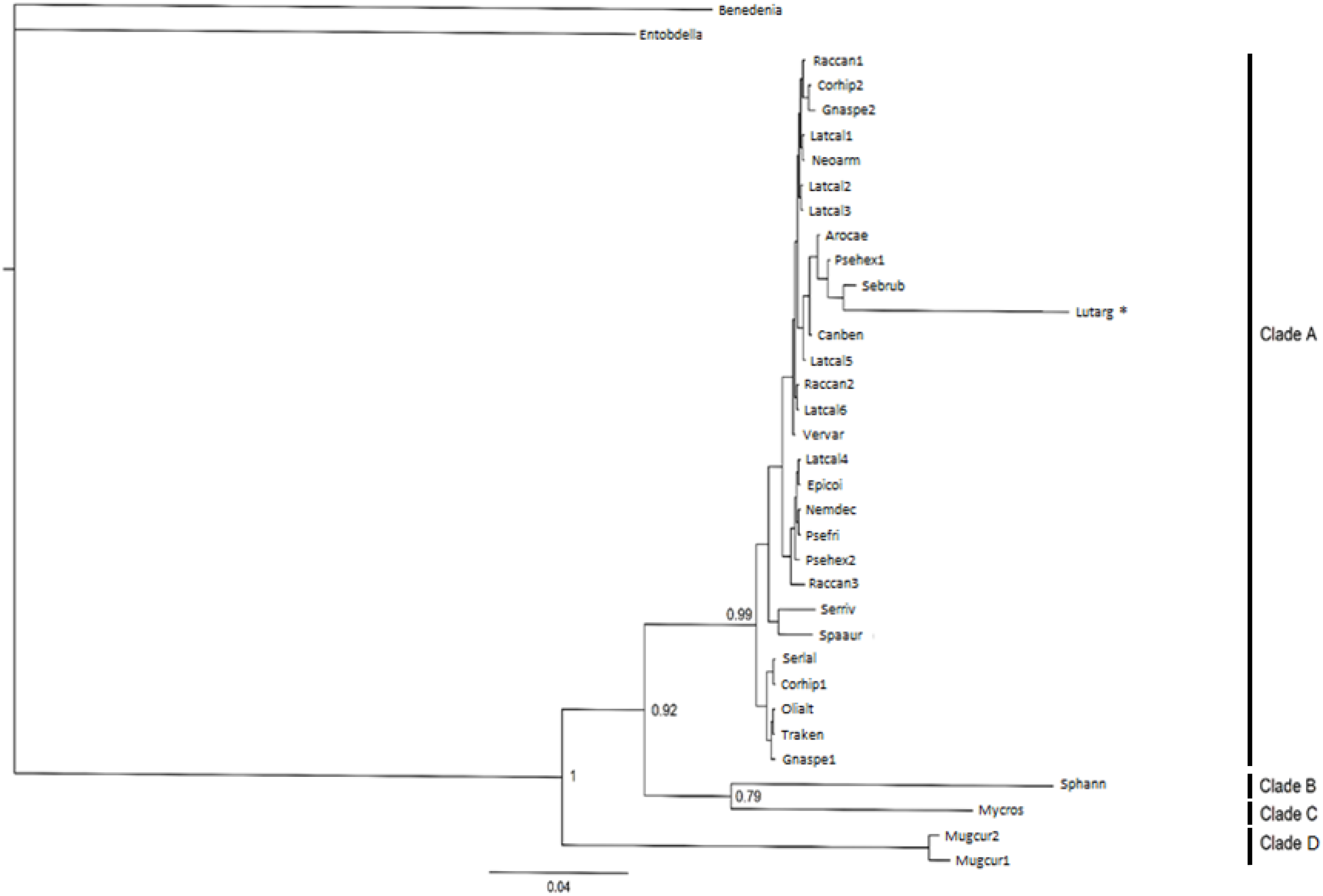
Relationships of species of *Neobenedenia* isolates collected from wild and captive host fish based on Bayesian inference and maximum likelihood analyses of the *H3, 28S rDNA*, and *cytochrome b* concatenated dataset. Posterior probability values are next to major nodes. “Benedenia” = *Benedenia seriolae* (HM222526.1 (*Seriola hippos* - South Australia, Australia), AY033941.1 (*Seriola quinqueradiata* - Japan), and FJ972088.1 (*Seriola hippos* - South Australia, Australia)) and “Entobdella” = *Entobdella soleae* (FJ972108.1 (*Solea solea* - United Kingdom), AY486152.1 (*Solea solea* - United Kingdom), HQ684799.1 (unknown host - China)) sequences obtained from Genbank were included as outgroups.

**Fig. 2.**
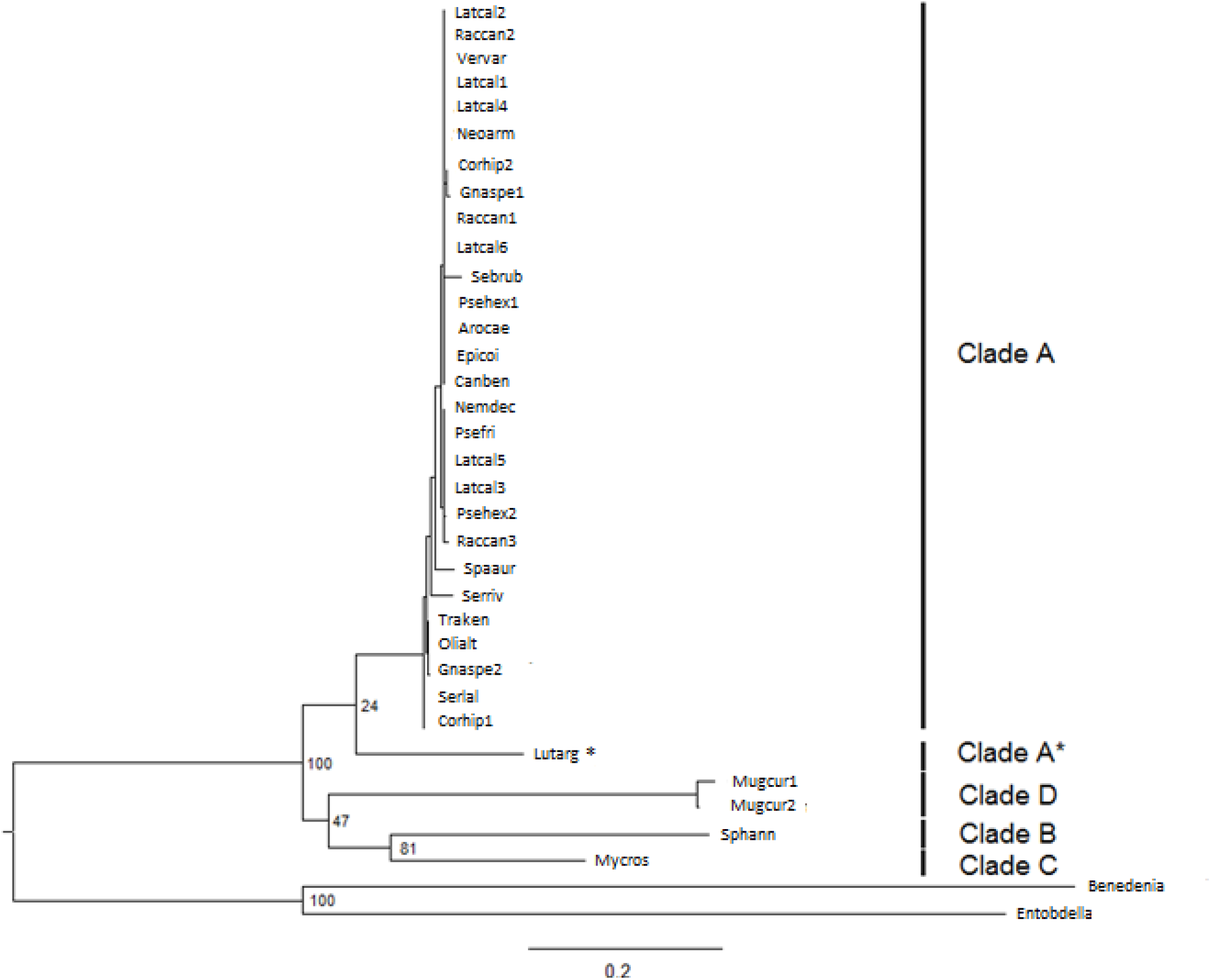
Relationships of species of *Neobenedenia* isolates collected from wild and captive host fish based on Maximum likelihood analysis of the *H3, 28S rDNA*, and *cytochrome b* concatenated dataset. Maximum likelihood bootstrap proportions are next to major nodes. “Benedenia” = *Benedenia seriolae* (HM222526.1 (*Seriola hippos* - South Australia, Australia), AY033941.1 (*Seriola quinqueradiata* - Japan), and FJ972088.1 (*Seriola hippos* - South Australia, Australia)) and “Entobdella” = *Entobdella soleae* (FJ972108.1 (*Solea solea* - United Kingdom), AY486152.1 (*Solea solea* - United Kingdom), HQ684799.1 (unknown host - China)) sequences obtained from Genbank were included as outgroups. Clades labelled as per the topology of Figure 1. * indicates the *Lutjanus argentiventris* sample from Clade A in Figure 1.

Samples that comprise clades C and D have been morphologically identified by us (IDW) as *N. longiprostata*, Bravo-Hollis 1971 and *N. pacifica*, Bravo-Hollis 1971, respectively, while samples from both A and B are morphologically ‘*N. melleni*’. Clade B contains a single specimen collected from *Sphoeroides annulatus*, while clade A contains 28 of the 33 in-group samples (excluding the sample from *L. argentiventris*) included in the study. The isolate currently in culture at the Marine Parasitology Laboratory at James Cook University, Townsville, Australia collected from *Lates calcarifer* (Bowen, Queensland) also fell into clade A

The neighbor-joining tree constructed from all available *Cytb* sequences is presented in Figure 3. Given the proportions of informative sites (Table 3 - Supplementary Material), it is not surprising that the topology is reflective of the Bayesian and ML analyses. Furthermore, the sample collected from *L. argentiventris* is recovered as the sister of the clade A in congruence with the ML analysis (Fig. 2; clade A*). All of the *Cytb* sequences downloaded from Genbank fall into clade A, except for accession HMM222533 collected from *S. annulatus* (Perkins et al. 2009), which falls into clade B.

**Fig. 3.**
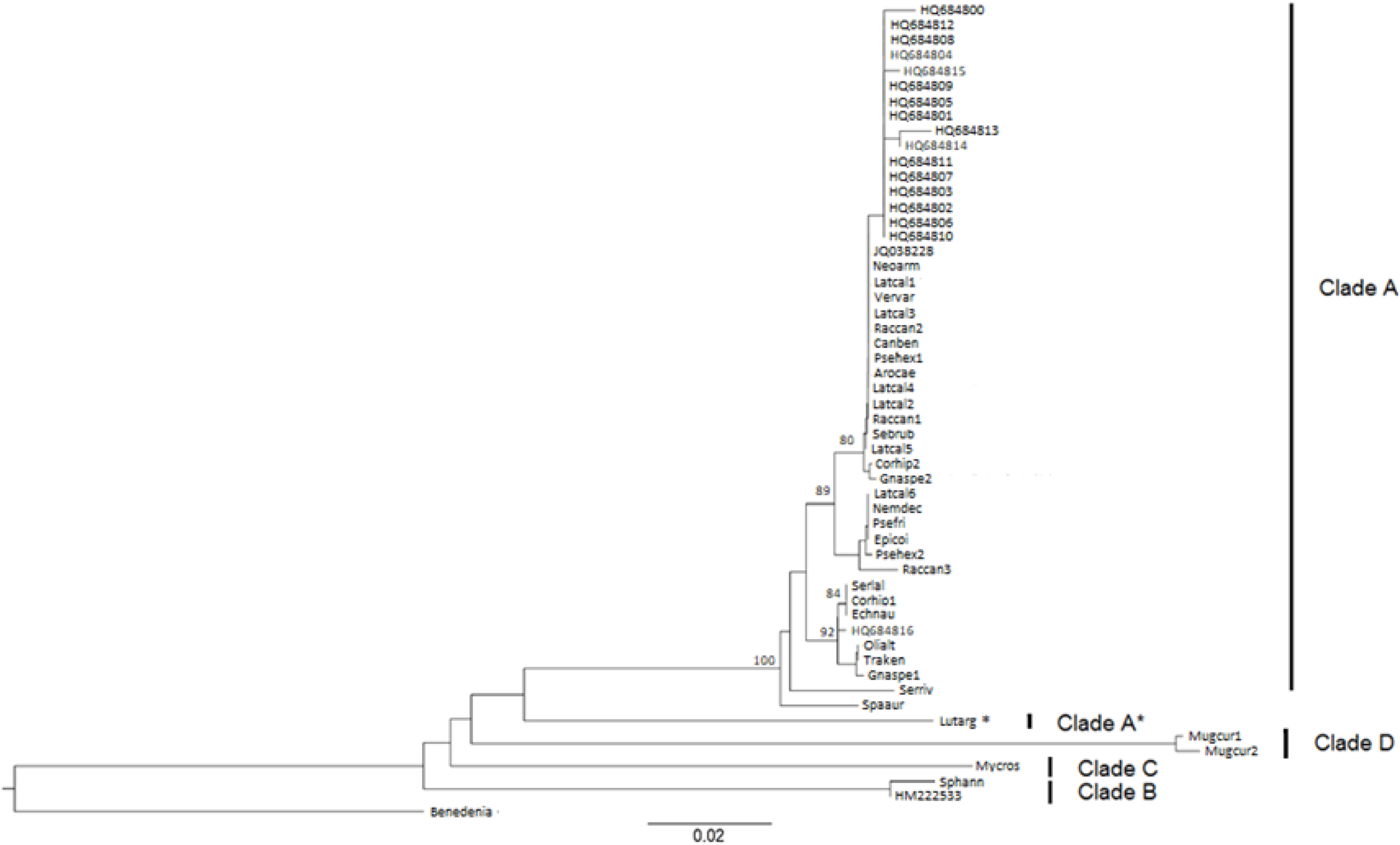
Relationships of species of *Neobenedenia* isolates collected from wild and captive host fish based on a Neighbor-joining analysis of partial *cytochrome b* sequences collected from Genbank and the present study. Genbank samples included in this analysis are titled using their Genbank identification code and were collected from a variety of host species. “Benedenia” = *Benedenia seriolae* (FJ972088.1 (*Seriola hippos* - South Australia, Australia)) sequence obtained from Genbank was included as an outgroup. Clades labelled as per the topology of Figure 1. * indicates the sample collected from *Lutjanus argentiventris* identified in Clade A in Figure 1.

Similarly, the neighbor-joining analysis (not shown) from all available *28S rRNA* sequences also recovered all previously identified clades. The specimen collected from *L. argentiventris* was once again recovered as a separate lineage but clusters with another specimen collected from *Sebastes rubrivinctus*, Jordan and Gilbert 1880, that was part of clade A in all other analyses. Several sequences fell into clade B but all these were sampled from *S. annulatus*.

## 4. Discussion

The correct identification of species underpins all biological study and for pathogens is critical in understanding infection dynamics, predicting outbreaks and determining effective control strategies. However, in the genus *Neobenedenia*, morphological plasticity, attributed to host induced morphological variation (Whittington and Horton, 1996), failure to accession specimens (including wet and mounted material) to curated collections (e.g. Cowell et al., 1993; Ellis and Watanabe, 1993; Koesharyani et al., 1999; Jahn and Kuhn, 1932; Müeller et al., 1992; Nigrelli and Breder, 1934; Nigrelli, 1935; Nigrelli, 1937; Nigrelli, 1947; Robinson et al., 1992; Zhang et al. 2014) and the lack of a robust genetic framework to aid in identification, has hampered these efforts. This study presents the most comprehensive phylogenetic investigation of *Neobenedenia* spp. to date, incorporating 33 isolates spanning 22 host fish species and nine countries and used both mitochondrial and nuclear gene markers.

Our results clearly show that there is a single species of *Neobenedenia* (Fig 1–3; clade A) that is both widespread geographically, able to infest a large number of different host fishes, and can be found on wild, farmed, and captive fish. Most importantly, clade A does not contain *Neobenedenia* specimens from *Sphoeroides annulatus*, which is likely to be the type host of *N. melleni* according to the description by McCallum (1927) and analysis by Whittington and Horton (1996). Our sample from *S. annulatus* (Fig 1–3; clade B), which was collected from wild fish in southern Mexico, is well differentiated from clade A and is the sister species to *N. longiprostata* in our analyses. Furthermore, except for HM222533, which was sampled from *S. annulatus* (Perkins et al., 2009) and clusters with our sample in clade B, all other *CytB* sequences obtained from Genbank which are named both *N. melleni* and *N. girellae* all fall in clade A, including the *CytB* sequence from the full mitochondrial genome (JQ038228), which is supposedly *N. melleni* (Zhang et al., 2014). Similarly, all Genbank *28S rRNA* sequences fall into clade A unless they were sampled from *S. annulatus*, where they clustered with clade B. Thus, the results of the present study support the findings by Whittington et al. (2004) and Perkins et al. (2009) that *N. melleni* and *N. girellae* are cryptic species in that they are morphologically very similar but genetically distinct. In an effort to stabilise the naming of *Neobenedenia* species, it seems prudent to ascribe *N. melleni* to individuals in clade B and retain *N. girellae* for those individuals in clade A.

It is likely that clade A* (Fig. 2, Fig. 3) represents another species of *Neobenedenia*, however at this stage, it is unclear whether this sample is a previously recorded *Neobenedenia* species or if it is novel to science. The single specimen represented in this clade is the only one collected from a lutjanid host (*Lutjanus argentiventris* collected from La Paz, Mexico) and the assessment of this family of hosts for potential *Neobenedenia* species is an important avenue of research to pursue. The Bayesian placement of this specimen into clade A appears incorrect given the long branch length and ML and neighbor-joining analyses clearly differentiate it from other samples in this study. Further work is underway to assess that status of this individual.

The findings of this study suggest that, historically, both prior and post the proposed synonymy by Whittington and Horton (1996), many of the parasites identified as *N. melleni* may in fact be *N. girellae* given how few isolates were identified as *N. melleni* in this study (e.g. Bullard et al., 2000; Deveney et al., 2001; Kerber et al., 2011; Landos, 2012; Wang et al., 2004; Zhang et al., 2014). Accurate identification of *Neobenedenia* species may be hampered by intraspecific morphological plasticity thereby resulting in false identifications. Despite the recognised difficulty in differentiating *N. melleni* and *N. girellae*, the majority of studies focussing on *Neobenedenia* spp. have not stated that they have accessioned samples to museum or private collections. This makes clarification of previous identifications almost impossible and accurate identification of historical host species and geographic locations for each of these two taxa unachievable. However, given the number of isolates that were included in this study and the varied host species and locations they were collected from, it is likely that the number of hosts and locations that *N. melleni* has been credited infecting and inhabiting has been overestimated. To facilitate morphological and genetic comparison of individuals in the future, it is recommended to accession reference material (both mounted and fixed specimens).

Using our genetic framework, it will be possible to reassess the morphological differences between *N. girellae* and *N. melleni a posteriori*. The *CytB* neighbor-joining analysis shows significant substructuring in *N. girellae*, which may account for the variable morphology noted by Whittington and Horton (1996). Morphological variation can be due to epigenetic factors which influences the phenotype expressed depending on the particular environment being experienced (Agrawal 2001; Mati et al., 2014; Olstad et al., 2009; Via et al., 1995). The flexibility displayed by these parasites makes the identification of species solely through morphological means considerably challenging (Barcak et al., 2014; Bickford et al., 2006). As closely related cryptic parasite species may exhibit variable traits to one another such as host susceptibility (Reversat et al., 1989), pathogenicity (Haque et al., 2003; Homan and Mank, 2001; Skovgaard, et al., 2002), and epidemiology (Murrell and Pozio, 2000), accurate identification is crucial to understand the risk posed by the presence of a particular species in a system.

*Neobenedenia girellae* infects a large number of fish species of economic importance including commercial fisheries, aquaculture, and the ornamental trade (Ogawa et al., 1995; 2006). *Neobenedenia girellae* isolates were collected from several fish species that support major aquaculture production or commercial fisheries including *Lates calcarifer* (barramundi or Asian sea bass), *Epinephelus coioides* (gold-spot grouper), *Coryphaena hippurus* (mahi mahi or dolphinfish), *Plectropomus leopardus* (coral trout), *Rachycentron canadum* (cobia), *Seriola lalandi* (yellowtail kingfish), and *Seriola rivoliana* (almaco jack), Valenciennes 1873 (Fig. 1). Similarly, attractive tropical species are also at risk. Parasites infecting eight species of popular ornamental fish species from seven families were among the isolates included in this study. All represent new host records for *N. girellae* and show the diversity of host species this parasite can infect (Table 1; see *3.1.)*. Ornamental fish support a huge, multi-national industry that involves hundreds of fish species (and their associated pathogen communities) being transported all around the globe (Bruckner, 2004). This provides an excellent opportunity for the dispersal of *N. girellae*, further encouraged by its lack of host specificity. Surveillance of this pathway for *N. girellae* is advised given its broad host specificity.

Isolates collected from two outbreaks on *L. calcarifer* which occurred in Queensland Australia, the first on a fish farm in Hinchinbrook in 2000 and the second in Gladstone in 2014 were included in this study (Table 1) and were determined to be a part of the *N. girellae* clade (Figure 1–2). The parasites of concern were initially identified as *N. melleni* by Deveney et al. (2001) and Landos (2012), however, our study suggests that these were erroneous identifications and the species associated with these mortality events was in fact *N. girellae*. Similarly, the previously unidentified species of *Neobenedenia* which is currently being cultured at the Marine Parasitology Laboratory at James Cook University, Townsville has also been identified as *N. girellae* using this analysis. As such, all previous research on this species from the Marine Parasitology Laboratory is now ascribed to *Neobenedenia girellae* (i.e. Brazenor and Hutson 2015; Dinh Hoai and Hutson 2014; Hutson et al. 2012; Militz et al. 2013a; 2013b; Militz and Hutson 2015; Trujillo-Gonzalez et al. 2015a,b).

The large diversity of host species that *N. girellae* is able to infect (23 host species determined from the present study) and the geographic range in which it is present (found between latitudes 23.8489° S and 24.1422° N (wild isolates) and 23.8426° S and 24.1426° N (all isolates) from samples included in this study) makes it a globally cosmopolitan species and a threat to aquaculture industries around the world.

## Acknowledgments

AKB was supported by an Australian Postgraduate Award and funding from the JCU Graduate Research School Research Grants program. Partial funding for this work was provided through a Queensland Government Smart Futures grant to TLM and KSH. This research was supported by use of the Nectar Research Cloud, a collaborative Australian research platform supported by the National Collaborative Research Infrastructure Strategy (NCRIS). Dr Leslie Chisholm, South Australian Museum and Dr Robert Adlard Queensland Museum accessioned specimens. We thank all colleagues who contributed specimens for this research: Roger Chong, Ariel Diamant, Mark Hilder, Noritaka Hirazawa, Thane Militz, Federico Rotman, David Vaughan, Erik Vis, and Roger Williams.

The authors would like to acknowledge Ian Whittington who passed away in October 2014. He was a greatly appreciated colleague and friend and without his enthusiasm of Capsalid research this study would not have been possible. He is deeply missed.

## Supplementary Material

**Table 1:**
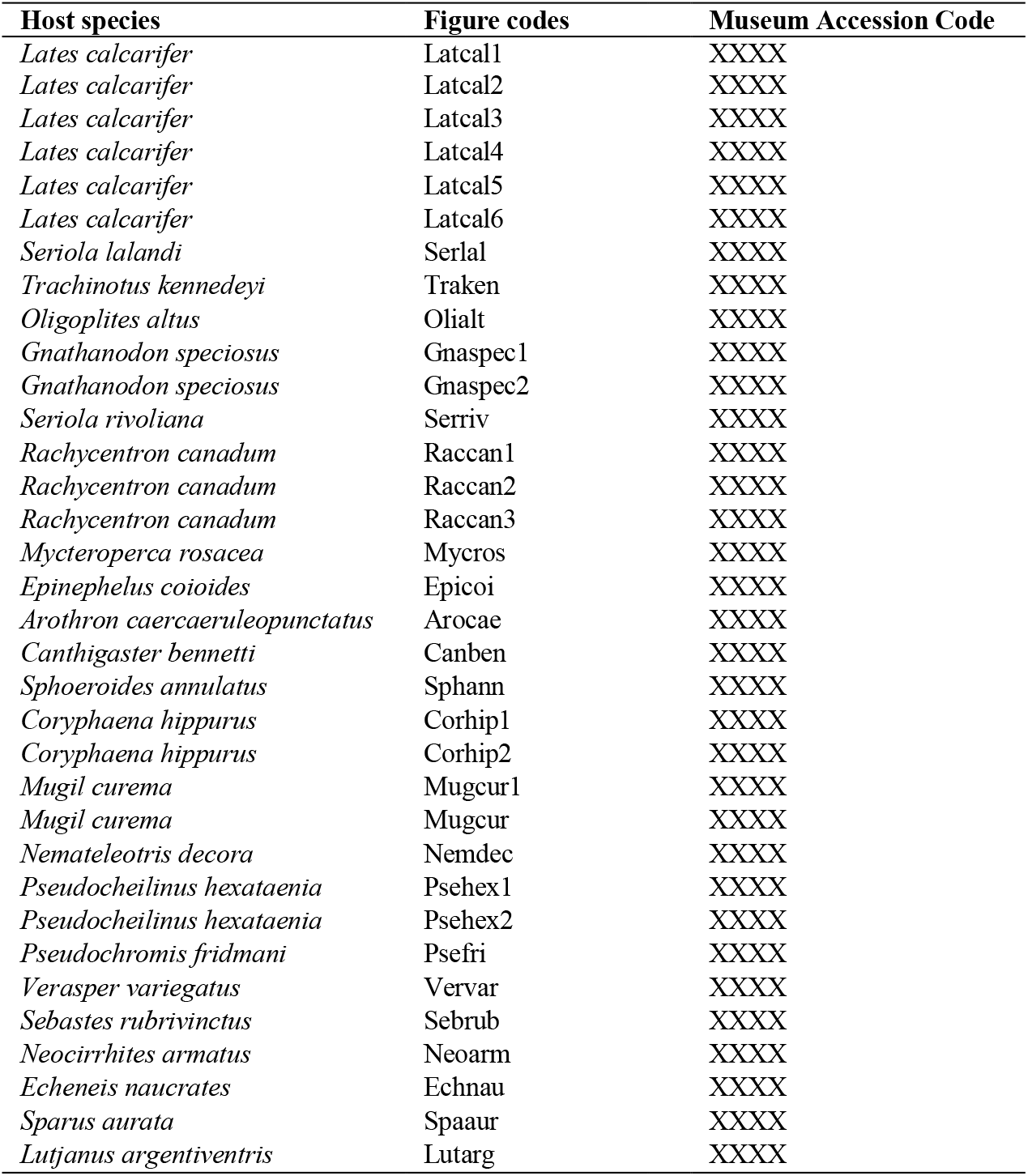
Museum accession numbers for *Neobenedenia* used in this study. XXXX = code forthcoming upon acceptance of the manuscript.

**Table 2:**
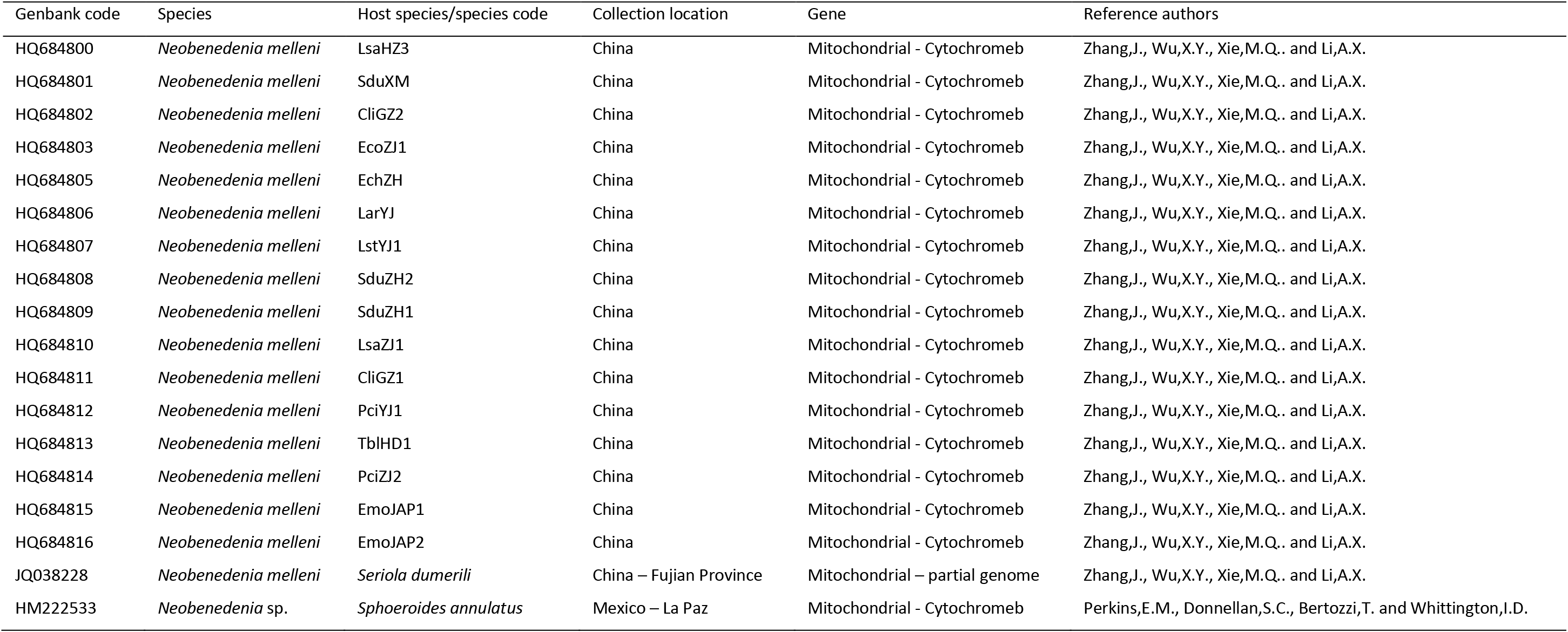
Sequences of *Neobenedenia* (identified as *“Neobenedenia melleni*” by researchers) retrieved from Genbank, the lodging researchers and associated codes

**Table 3:** Proportion of informative sites of three genes; 28S, H3, and cytochrome b for sequences included in analyses.

| Gene | Length | Parsimony informative sites |
| --- | --- | --- |
| 28S | 411 | 24 |
| H3 | 292 | 26 |
| Cytochrome b | 704 | 166 |

